# Genome-wide identification of intrasplicing events in the human transcriptome and hints to their regulatory potential

**DOI:** 10.1101/159350

**Authors:** Maximilian Radtke, Ismet Srndic, Renée Schroeder

**Affiliations:** Department of Biochemistry and Cell Biology, Max F. Perutz Laboratories, University of Vienna, Dr. Bohr-Gasse 9/5, 1030 Vienna, Austria

**Keywords:** Intrasplicing, human transcriptome, nested splicing, recursive splicing

## Abstract

Alternative splicing is one of the major regulators of both, transcriptome diversity and individual isoform abundance. Therefore regulation of alternative splicing is crucial and yet, due to the complexity of the human genome (23.000 genes, most of which can be alternatively spliced) diverse and multileveled. Identifying and understanding the scope and variety of splicing events is still an ongoing process. We established a novel pipeline to extract splicing events from specific RNA-sequencing datasets and identified numerous splicing events that did not span the entirety of the respective annotated intron but used intronic splice sites. These splicing events could be generally categorized into three groups: 5’recursive (using the exonic splice donor and intronic acceptor), 3’recursive (using an intronic splice donor and an exonic acceptor) and nested (using two intronic splice sites). Surprisingly, the splicing events we found occurred in introns of all lengths, but generally followed the abundance scheme of all introns, i.e. most were found in introns between 500 and 5000 bps. After confirmation of these splice events by conventional methodologies, we further analyzed the impact of intrasplicing on full intron removal. For this we established a luciferase-based reporter which showed that these intronic splicing steps can be beneficial, deleterious or neutral for full intron removal. Thus intrasplicing events can be crucial for determining the transcriptional output. This in part confirms recent findings on recursive splicing events in humans and other vertebrates and further uncovers an additional level of transcriptome regulation based on a yet undiscovered level of flexibility and regulation of splice site selection and its impact on gene expression.

## Introduction

Precursor messenger RNA (pre-mRNA) splicing is not only a key step in protein production, but is also critical for the regulation and expansion of the functional proteome. The human genome contains an unexpectedly low number of genes, approximately 23,000 and alternative splicing in combination with posttranslational modifications is known to diversify these into 90,000 proteins^1^. High-throughput sequencing studies suggest that up to 100% of human multi-exon genes produce at least two alternative mRNA isoforms. The majority of human introns contain 5’- and 3’-splice sites with a consensus sequence of variable conservation providing room for many ways how to define a splice site^2^. Combinatorial events including RNA-protein interactions and RNA-RNA interactions have been shown to be essential for determining splicing events, especially in long introns. The longer the intron, the higher the precision needed to control splice site selection.

### Mechanisms of long intron splicing

Different mechanisms of long intron splicing in higher eukaryotes have been proposed over the last two decades, with two main models emerging: recursive and nested splicing^3–6^ (Figure 1). Nested splicing is an intron-internal splicing event that does not use the primary or exonic splice sites. Whether or not this type of splicing is a necessity for effective splicing of a subset of (longer) introns is still unsolved. Splice site-like sequences, dispersed over introns are able to prevent premature cleavage and polyadenylation (PCPA)^7^ by binding U1snRNPs and could promote intra-splice site usage. In a single case, the regulatory impact of intrasplicing has been demonstrated: in the 4.1R gene alternative promoter usage in combination with subsequent and first-exon-dependent intrasplicing determines the expressed isoforms^8^.

**Figure 1:**
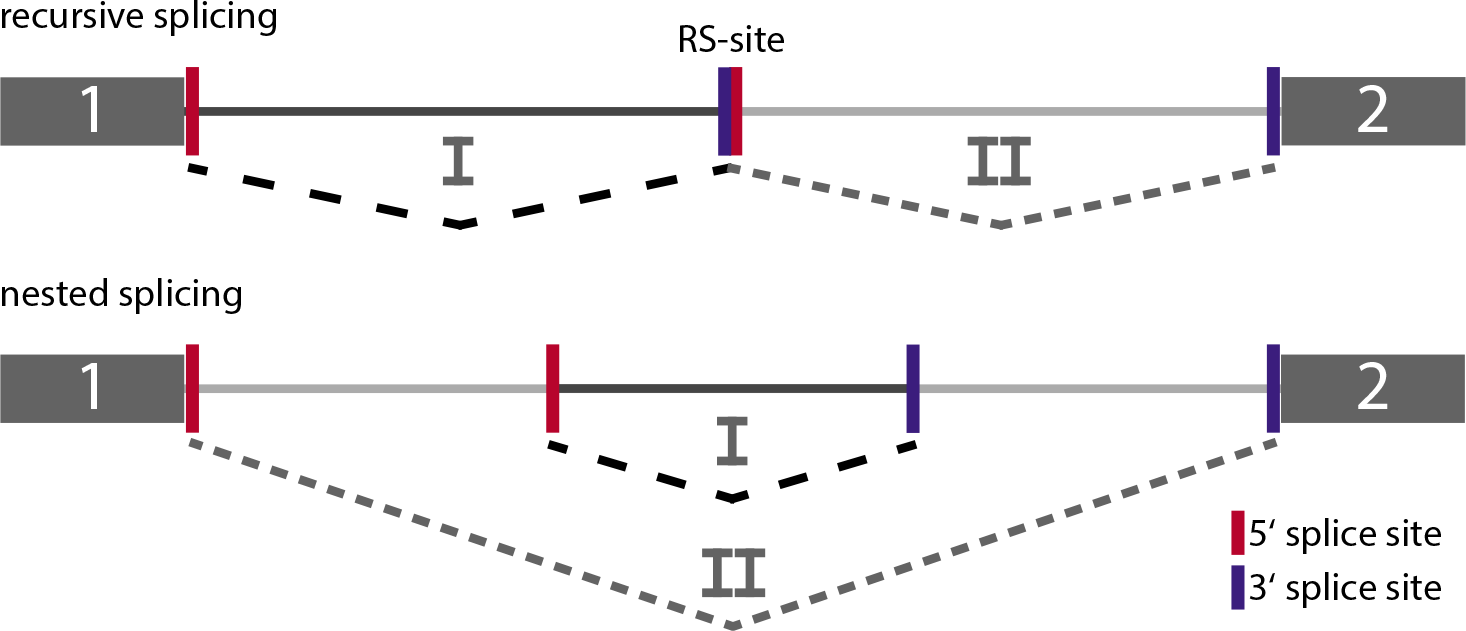
Proposed splicing models for long intron splicing. Recursive splicing (top) uses intronic recursive splice sites (RS-sites) that reconstitute a new 5’ or 3’ splice site after a first splicing step, depending on the directionality of the recursive splicing cascade. Nested splicing (bottom) uses two intronic splice sites that remove part of the intron before the final splicing step removes the entire intron.

Recursive splicing (RS), in contrast to intrasplicing, involves incremental splicing reactions. One of the primary splice sites is involved and intronic splicing reconstitutes a functional, recursive splice site (RS-site) that is utilized for subsequent splicing steps (Figure 1A). Two recent studies analyzed recursive splicing in depth and on large scale in drosophila and vertebrates, demonstrating that this mechanism is prevalent in the longer introns. In humans, RS does not seem to be a necessity for long intron removal but might rather have a regulatory impact on in- or exclusion of RS-exons^9^. And as shown in D. *melanogaster*, intron length cannot be the only requirement to induce recursive splicing cascades^10^, making space for regulatory potential of these splicing events.

### Detection of splicing events

One of the major obstacles in detecting and verifying intermediate splicing events such as nested and recursive splicing, is the temporary nature of their products. Partially spliced or pre-spliced pre-mRNA is subjected to immediate further processing (i.e. subsequent splicing steps). Lariats arising from recursive or intrasplicing events are, presumably as most lariats, turned over by debranching with Dbr1 and exonucleolytic degradation of the intron RNA within minutes after splicing^11,12^. A combination of PCR-based methods, lariat enrichment and deep sequencing can be applied to partially overcome these restrictions^13,14^. Lariats can also be stabilized by interfering with the degradation pathway, mainly by inhibiting debranching. Lariat sequencing as performed by Awan et al.(2013)^14^ took advantage of a Δ*dbr1* strain, resulting in the identification of hundreds of novel splicing events in *S. pombe*. In more complex genomes which require a more stringent splicing regulation due to the increased abundance of potential alternative splicing events and longer host introns, we found that Dbr1 knockdown to non-lethal levels (∼40% of wildtype) resulted in a major shift in genome wide splicing patterns and introduction of new, aberrant splicing events (unpublished). Therefore this approach is not applicable, at least in human cells.

Another restriction in lariat identification, if approached with RNA-sequencing, is an increased occurrence of mapping artifacts due to repetitive elements within introns and general sequence similarities at splicing regulatory sequence elements. These artifacts deteriorate when reads spanning either exon-exon or branchpoint-5’ss junctions (split-reads) are required to identify lariats and splice sites, reducing the effective mapping length (Figure 2).

**Figure 2:**
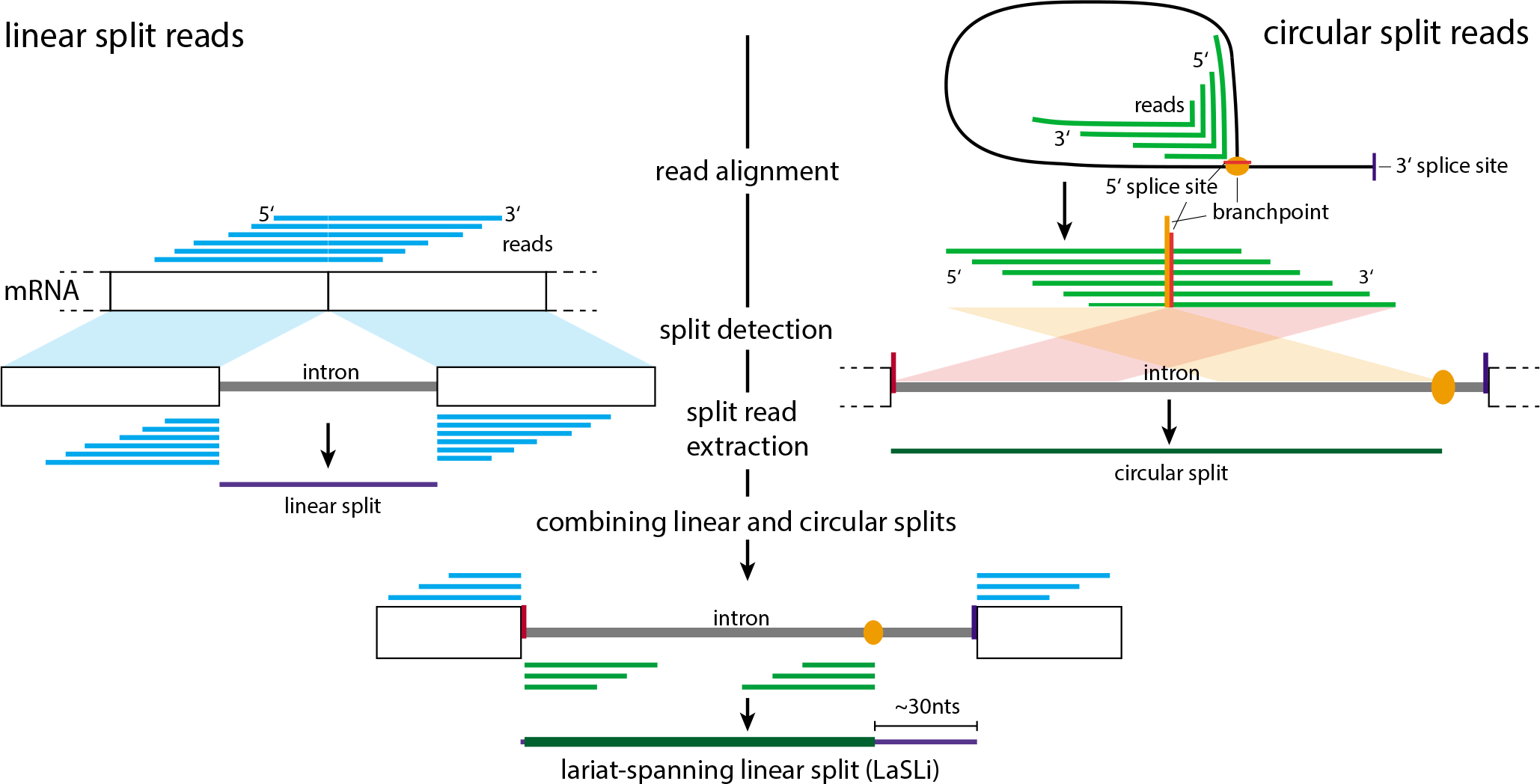
Split-reads that cover exon-exon or branchpoint-5’splice site junctions are extracted from segemehl-aligned RNA-seq datasets. Split reads originating from the same splicing event (exon-exon junction reads and lariat reads) can be combined in order to obtain splice site and branchpoint positions (see Materials and Methods). This allows for a higher confidence mapping of splice sites than previous approaches which rely solely on linear split reads.

Previous studies^15–18^, dealing with the detection of branchpoints^16^ or novel splicing events strongly rely on splice site annotations, which allows an event prediction with very high precision, but does not aim at the detection of novel, intronic splicing events. A promising approach in this direction is targeted RNA-seq with extremely high coverage and subsequent analysis by SplicePie^17^ that allows to draw a wealth of conclusions about splicing sequentiality, unusual splicing events and intron dwell times. Yet, this analysis focuses on single genes to achieve the required high read coverage and does not allow for a genome wide analysis.

### Lariat Spanning Linear Split Reads

In this work, we performed an in depth analysis of existing RNA-sequencing datasets of human cells using detailed bioinformatic analyses to identify unusual splicing events in a genome wide manner. To overcome the above described restrictions, we applied an approach that makes use of the vast quantity of published RNA-seq datasets, focusing on those generated with a library preparation protocol that enriches RNA species that are likely to hold information on unusual splicing events. The novelty of this procedure is the combination of reads covering the linear splice junction with reads that arise from the branchpoint-5’ splice site junction of lariats (Figure 2). These **la**riat **s**panning **li**near **s**plit reads (LaSLis) therefore represent a set of high confidence splicing predictions. This approach resulted in the identification of more than 90,000 splicing events (neglecting alternative branch points), of which 5,693 are not covering the full length of annotated introns and are therefore potential recursive or intrasplicing events. As recursive splicing in the context of long introns has already been thoroughly investigated^5,9,10^, we focused on those events found in shorter introns (<5000 nucleotides), which comprised the majority of the extracted splicing events (see Figure 3A). The logic behind the model of recursive splicing as a means for facilitating long intron splicing excludes the necessity of such events for short intron splicing. Therefore, these splicing events could hold regulatory capacity.

**Figure 3:**
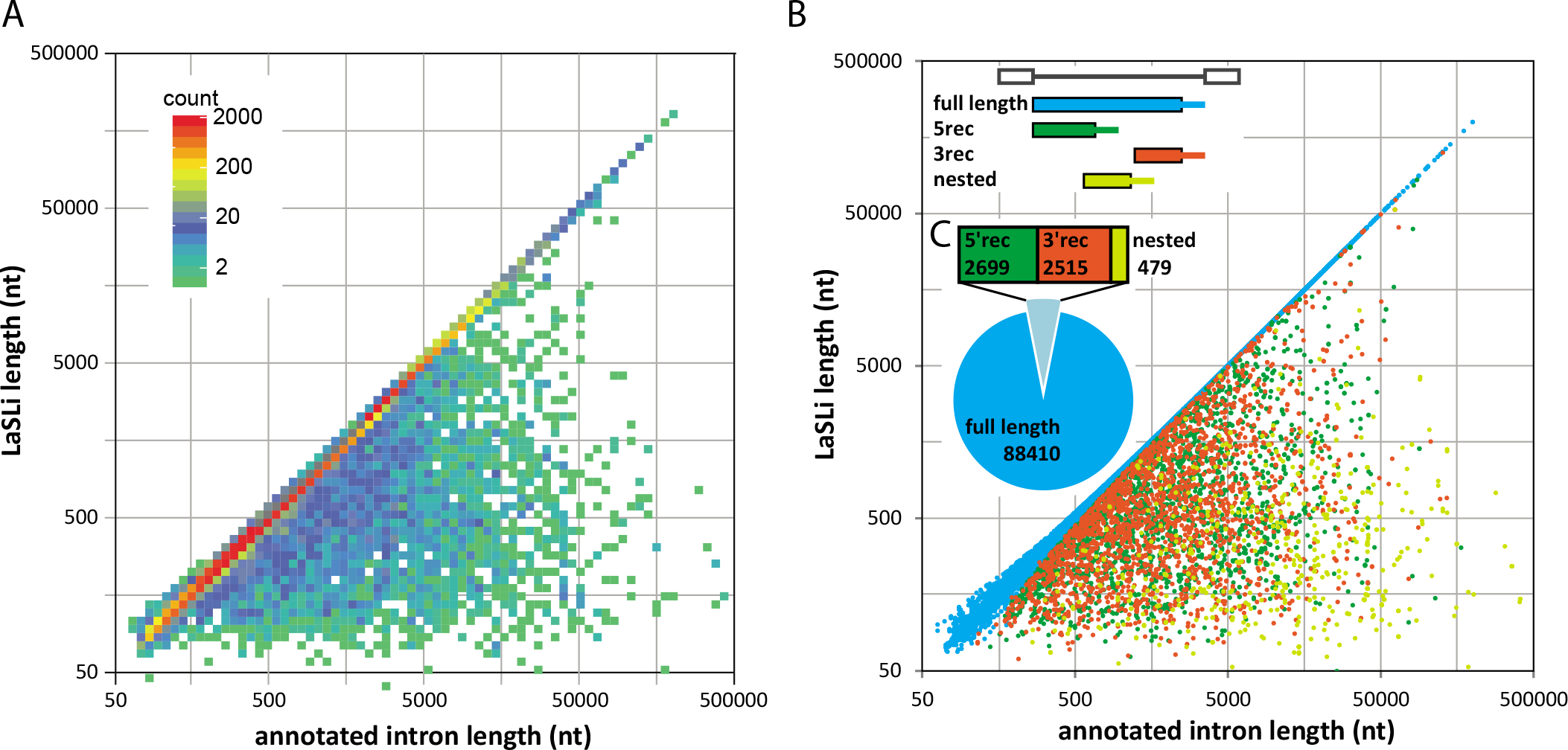
Overview of the extracted splicing events. (A) shows the length of the extracted splicing event plotted against the length of the corresponding intron. This plot is shown as a density plot to allow for visualization of overlapping splicing events, grading the number of splicing events with similar properties by the colour grade shown in the legend. The majority of splicing events, both, full length and shorter are found in introns < 5000 nts. In (B) the same splicing events are classified based on their position in the corresponding intron: full length (blue), putative initial 5’ recursive (green), 3’ recursive (red) and nested splicing events (yellow). The abundance of each class of splicing event is shown in (C).

## Results

### Extraction of novel splicing events

The aim of this work was to identify rare and non-canonical splicing events in the human transcriptome and to test if these events affect splicing output, which would point to a potential regulatory function. To achieve this, we established a novel extraction pipeline, designed to extract transient splicing events with high confidence. This approach utilizes reads from RNA sequencing datasets that cover both, linear splice junctions and circular branchpoint junctions (Figure 2). The combination of these two split-read types under certain splicing-specific criteria such as branchpoint distance and splice site positioning (see Methods), allows for stringent filtering and strongly reduces mapping artifacts that accompany split-read alignment. This approach additionally maximizes the alignment length and thus allows for the identification of novel splicing events, outside the known intron-exon boundaries.

The datasets we mainly focused our analysis on were generated by library preparation protocols that enriched either for long non-polyadenylated nuclear RNA^19^, by size selection and fractionation, or circular RNA by exonuclease treatment or CaptureSeq^16,20^. The former includes, besides other RNA species, precursor- and partially processed mRNA, which sets the foundation for extraction of linear splice junctions. The latter holds, besides circRNAs, reads of lariats, which are the basis for branchpoint split read extraction.

### Splicing event classification

Using this approach we identified 97,338 splicing events (neglecting alternative branchpoints), most of which span full introns. 5,693 events are shorter than the respective annotated intron and therefore may originate from recursive or nested splicing. The majority of non-full length splicing events are found in introns shorter than 5000 basepairs (bp) (Figure 3A). Further classification of this dataset subdivided the splicing events into 4 classes:

1) Full length (**FL**): covering a fully annotated intron, allowing for alternative splice sites within 50 nucleotides of the annotated splice site (88,410 events).
2) 5’ recursive (**5rec**): the splicing event uses an annotated 5’ splice site (or one close to it) but does not extend throughout the entire intron, using an unannotated, intronic 3’ splice site, thus being a potential first splicing event of a recursive splicing cascade (2,699).
3) 3’ recursive (**3rec**): like 5rec, with the difference, that the 5’ splice site is within the intron and the 3’ splice site is annotated (2,515).
4) Nested splicing event (**nest**): both, the 5’ and the 3’ splice sites lie within the intron and neither is annotated (479).

The distribution of all splicing events identified over these four classes is depicted in Figure 3B and C. These numbers do not take into account alternative branchpoints, but show only those events with distinct 5’ and 3’ splice sites.

In order to keep naming of different types of splicing events consistent, in this paper we use the following designations: **FL**, **5rec**, **3rec** and **nested** as described above; **intrasplicing** refers to a splicing event that does not span the entire annotated intron and can therefore be a recursive or nested splice; **LaSLi** stands for Lariat Spanning Linear split and describes splicing events identified in our extraction pipeline; Splice sites that do not mark an annotated exon-intron border are termed **intra-splice sites**.

### Data availability

The full dataset, including splicing event classification, splice site scoring and alternative branchpoint positioning is available in the supplementary materials. Note that circular split read positions, i.e. potential branchpoint positions, are only available in the set *all_laslis_BP_bed8.bed* as block coordinates. Splice site scoring is available as a separate set of files with the name column (4) of the bed format as the 5’ss score and the score column (5) as 3’ss score.

### Characteristics of intronic splicing events

In order to gain an overview about the splice site quality of LaSLis, we determined the splice site strength of splice sites of each splicing class with Xmaxentscan^21^, a tool that scores a given sequence by the maximum entropy principle and references to a predetermined splice site consensus (http://genes.mit.edu/burgelab/maxent/Xmaxentscan_scoreseq.html).

The score distributions for each intrasplicing class as well as annotated splice sites are given in Figure S 1. Average scores for annotated splice sites are 7.84 for 5’ and 7.99 for 3’ss. 5rec and 3rec exonic sites are within that range, with the intronic splice site scoring at 4.31 and 5.70, respectively. Both intra-splice sites of nested LaSLis have a much more spread out score distribution with a strong difference between 5’ and 3’ splice sites (Figure S 1C). We used these scores, to adjust splice site definition and overcome the error prone split-read mapping. In this approach we allowed for a maximum adjustment of 2 nucleotides at each splice site and selected the highest scoring position. Especially 3’splice site assignment benefitted from this approach and resulted in a final dataset with much higher splice site assignment fidelity.

Next, we performed a meta-analysis on these 4 classes of splicing events to investigate and compare their properties, with respect to annotated splicing sites. This also serves as a measure of the performance of our extraction pipeline as the unique properties of introns and lariats, such as branchpoint and splice site consensus sequence, branchpoint – 3’ss distance, conservation, etc., are well studied and can be referenced to. The vast majority of splicing events that are in contact with an exon, such as full length, 5rec and 3rec, fit the annotation in terms of splice site position (Figure 4A). Interestingly, those events that show an offset to the annotated splice site follow a pattern of peaks every third nucleotide up- and downstream of the annotated splice site. This is indicative of alternative splice sites, which do not break the reading frame. Notably, alternative 3’ss seem to have a strong preference for a shift of -12 nucleotides into the intron, leading to additional 4 amino acids included upstream of that exon.

**Figure 4:**
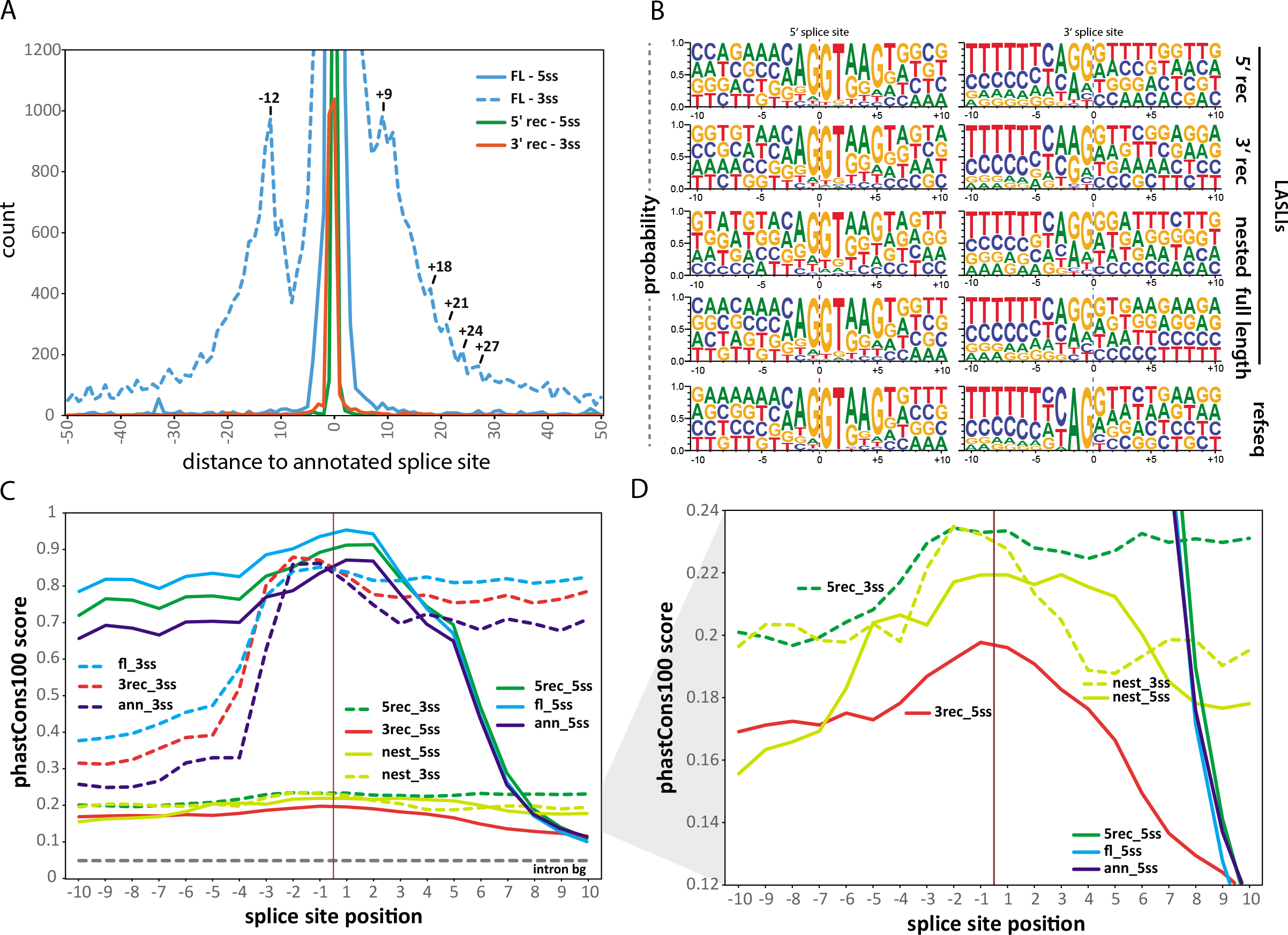
Splice site analysis of the 4 classes of splicing events. (A) shows the distance of the newly identified splice sites to the closest annotated splice site. (B) Splice site motifs of the four splicing classes as compared to the reference motif of 10,000 annotated introns. (C) shows the phastCons100 score across the splice sites. Intronic splice sites are generally less conserved and yet show a specific increase in conservation just around the splice site, as can be seen in (D), which is a close up of the indicated area of (C).

We next analyzed the sequences surrounding the splice sites to resolve any deviating splice site motifs for intronic splicing events (Figure 4B). Full length, 5rec and 3rec splice sites showed a strong GT-AG motif, very close to that of annotated splice sites, accounting for the precision of splice event extraction. Nested splicing events deviated strongly from the consensus sequence and reevaluation of the mapping data revealed that a lot of sequences originating from ribosomal RNA were falsely mapped to intronic regions. The abundance of these sequences allowed for a number of reads to evade our filtering criteria and get included in the nested splicing event subset. To remove these false positives, nested splicing events were further filtered by presence of a splice site consensus, i.e. GT at the 5’ and AG at the 3’ end. We have preliminary results, that also showed other, non-GT-AG nested splicing events to be spliced out, but these are subject to further studies and are therefore not included in the present dataset.

The conservation of splice sites was determined by extracting the corresponding genomic positions from the phastCons100 dataset, that calculates element conservation scores for 100 vertebrates^22,23^. Conservation scores of splice sites at intron exon boundaries show an expected drop in conservation towards the intronic region (Figure 4C). Splice sites located within the intron such as 3’ splice sites of 5’ recursive, 5’splice sites of 3’ recursive or both splice sites for nested events have a generally lower conservation, which still exceeds that of intron background and show, for the putative recursive splice sites, a similar conservation pattern as exonic splice sites. The conservation scores drop towards the site of the intrasplicing event (Figure 4D), which could be indicative of cryptic or RS exons. In contrast, nested splice sites display an increased conservation for approximately 10 and 6 nucleotides surrounding the 5’ and 3’ splice site, respectively.

The next characteristic we took a closer look at was branchpoint properties. The distance between the branchpoint and the downstream 3’ splice site is limited due to the structure of the spliceosome^24^ and can be used to determine the quality of our extracted splicing events. As shown in Figure 5A, the branchpoint distance peaks between 20 and 30 nucleotides, which concurs with literature^25^. Also the branchpoint sequences mainly follow what is known from literature, namely CUNAN or UUNAN^16^. Nested splicing events deviate from displaying a strong branchpoint adenosine. Deviations from consensus are likely to be introduced by the imprecision of split read mapping. The polypyrimidine tract downstream of the branchpoints is apparent for all splicing event classes (Figure 5C).

**Figure 5:**
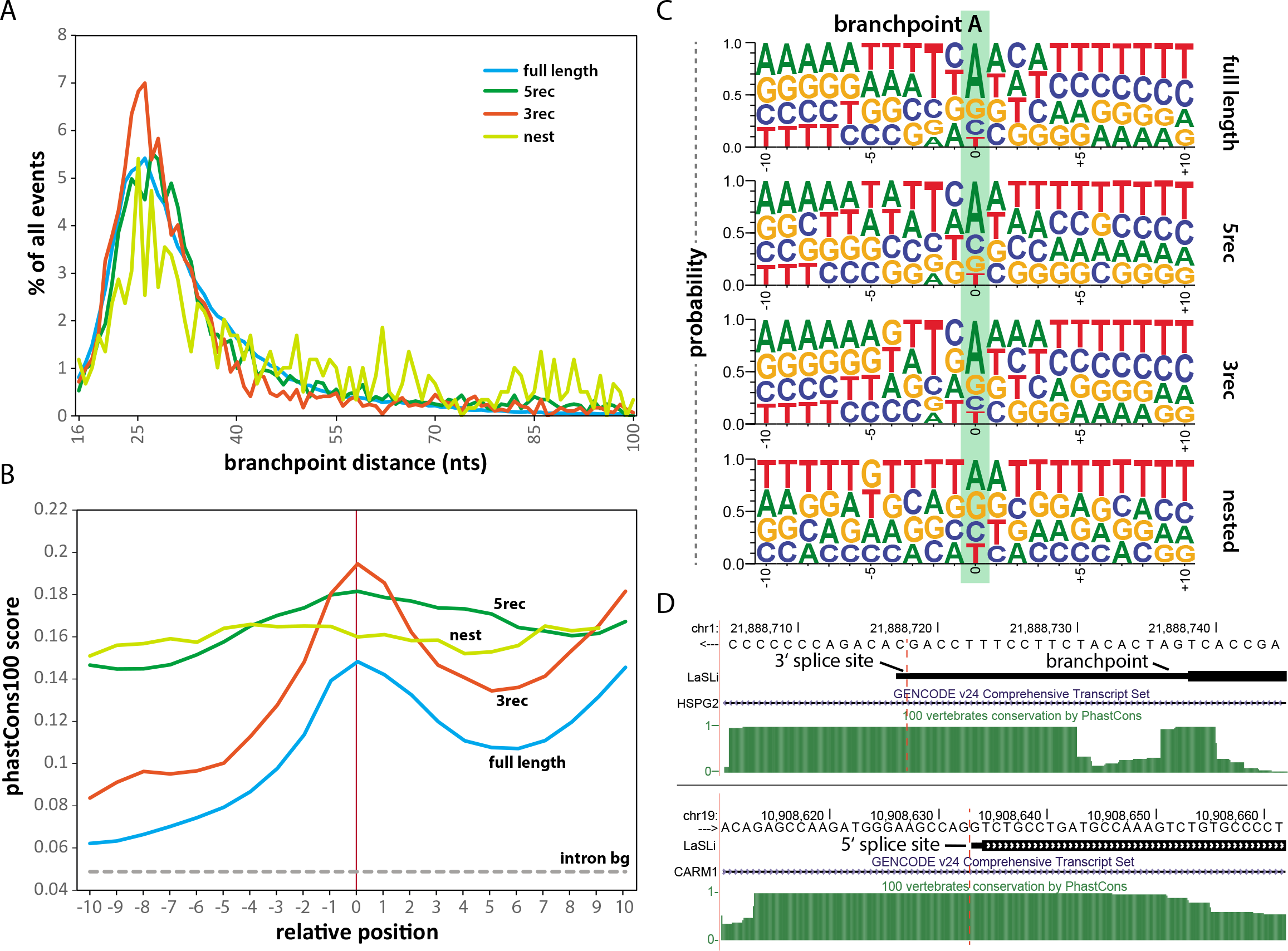
Analysis of branchpoint properties. (A) distance between the branchpoint and the respective 3’ splice site plotted against the fraction of events identified in that respective class. (B) branch point motif of the 4 LaSLi classes. (C) branchpoint conservation with phastCons100 score. (D) shows an example of a highly conserved 3’ splice site and branchpoint of a 5rec LaSLi in the *HSPG2*gene and a 5’ splice site of a 3rec LaSLi in *CARM1*. The bold bar indicates the circular, the thinner bar the linear split junction, corresponding to branchpoint and 3’ or 5’ splice site, respectively.

The conservation pattern at branchpoints shows an increased score for the branchpoint A and flanking nucleotides as compared to intron or local background (Figure 5B). Nested and 5’ recursive splicing events, and therefore branchpoints of exon free, intronic 3’ splice sites, have a similarly high score, yet do not show the characteristic drop in conservation before and after the branchpoint. Examples of a highly conserved branchpoint and intra-splice site are depicted in Figure 5D.

### Confirmation of unusual splicing events by lariatPCR

In order to confirm the existence of the computationally extracted splicing events, we performed lariatPCR on cDNA after RNA extraction from human cells and reverse transcription using random primers. This approach uses diverging primer pairs, binding downstream of the 5’ splice site and upstream of the branchpoint so that a PCR product is only obtained if a circularization event, i.e. lariat formation, has occurred (Figure 6A). LariatPCR further allows to precisely determine the site of the splicing event. We were able to obtain PCR products for the majority of lariats tested (Figure 6B). In many cases nested PCR was required due to the low abundance of lariats in the cells. RNA from HeLa, Hek293, K562 and Hep2G cells was used to increase the chances of detecting the lariats. PCR products were then sanger-sequenced and mapped against the genome to confirm the nature of the lariat and control for the precision of the splicing event extraction pipeline (Figure 6C). In case of the putative 3’recursive splicing event in *prkab2*, intron 7, we could also detect the linear splice junction on the preprocessed pre-mRNA (Figure 6D-F). The transient nature of lariats and splicing intermediates makes this confirmation process tedious, as even annotated, full length introns of highly expressed genes could not always be detected, presumably due to sequence repetitiveness and quick turnover. These properties inherit to and deteriorate in rarer or less abundant (due to low gene expression) splicing events. Thus our rather high confirmation rate (10/14) demonstrates the fidelity and robustness of our splicing event extraction approach.

**Figure 6:**
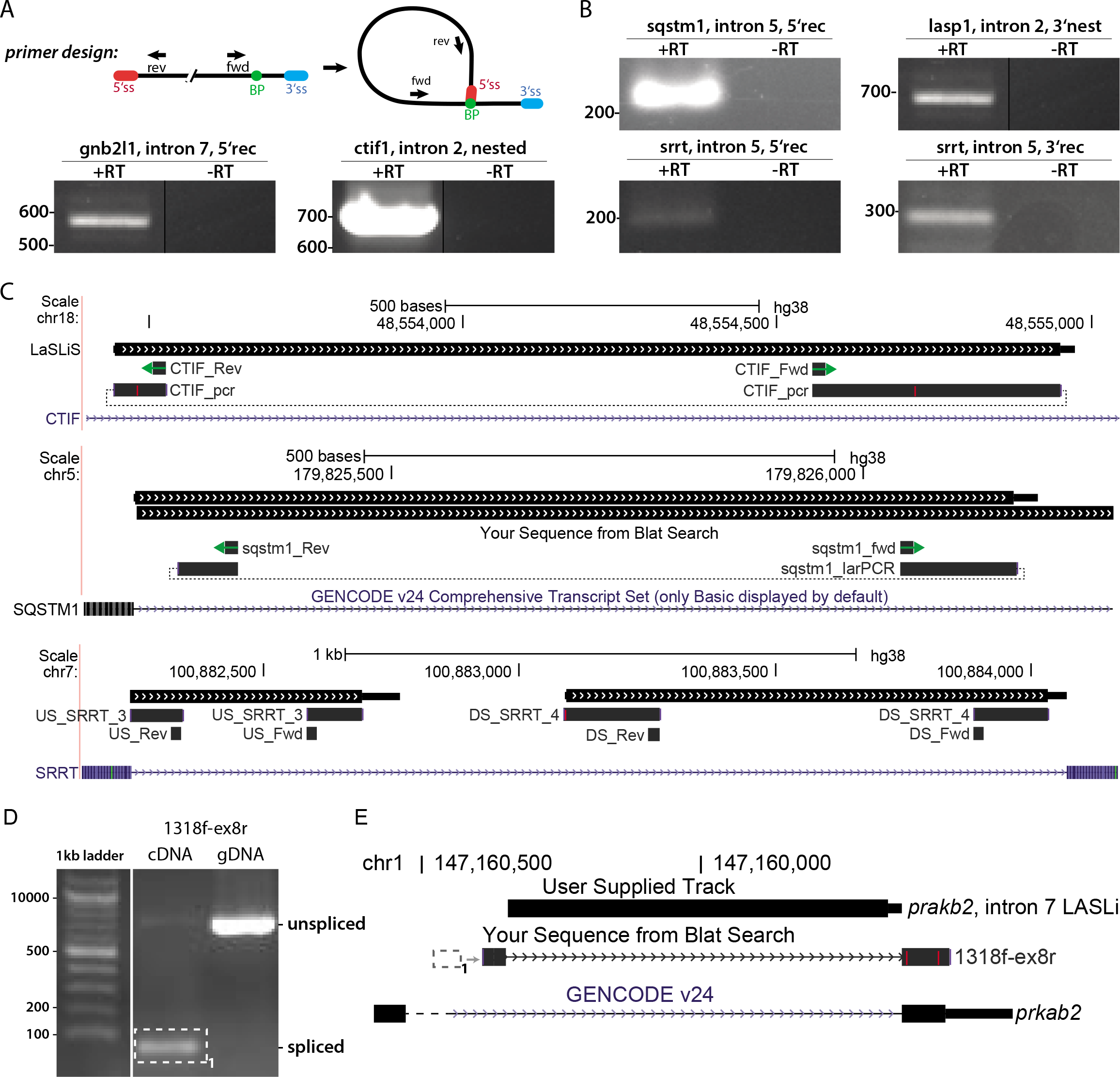
LariatPCR confirms the computationally extracted splicing events. (A) in order to obtain lariat specific PCR products, diverging primers were designed. Only in the case of a circularization during splicing and lariat formation, a PCR product will be obtained. (B) lariat specific PCR products obtained in a nested PCR. (C) larPCR products were sanger-sequenced, the obtained sequences were aligned to the human genome with BLAT and compared to the LaSLi dataset. Green arrows indicate the primer positions used for the larPCR. In few cases, linear splice junctions of nested splicing events could also be detected. (D) shows the primer positions in intron 7 and exon 8 of *prkab2*, (E) shows the pcr product obtained over the splice junction, compared to genomic DNA and (E) visualizes the mapping of the sequenced PCR product.

### The impact of intrasplicing on full intron removal in a reporter assay

To investigate the impact of intronic splicing events on full intron splicing efficiency and potentially transcript fate, we constructed a splicing reporter using the renilla luciferase, on the basis of the phRL-TK vector (Promega). Small introns (< 5000nts) were selected that showed at least one putative nested or recursive splicing event in our dataset, and cloned into the renilla gene at a position that favors splice site recognition (see Methods). The intra- or recursive splice sites were mutated or, in case of a 3’splice site, deleted together with the upstream polypyrimidine tract. Further in depth analysis was performed by removing the nested splicing event on DNA level, elucidating the fate of the pre-spliced transcripts. Vice versa, we also generated constructs, containing only the nested splicing event to evaluate their splicing efficiency, allowing us to draw conclusions about intra-splice site strengths (see Figure 7A). Using this approach, we identified three different types of effects of intrasplicing events on full intron splicing.

**Figure 7:**
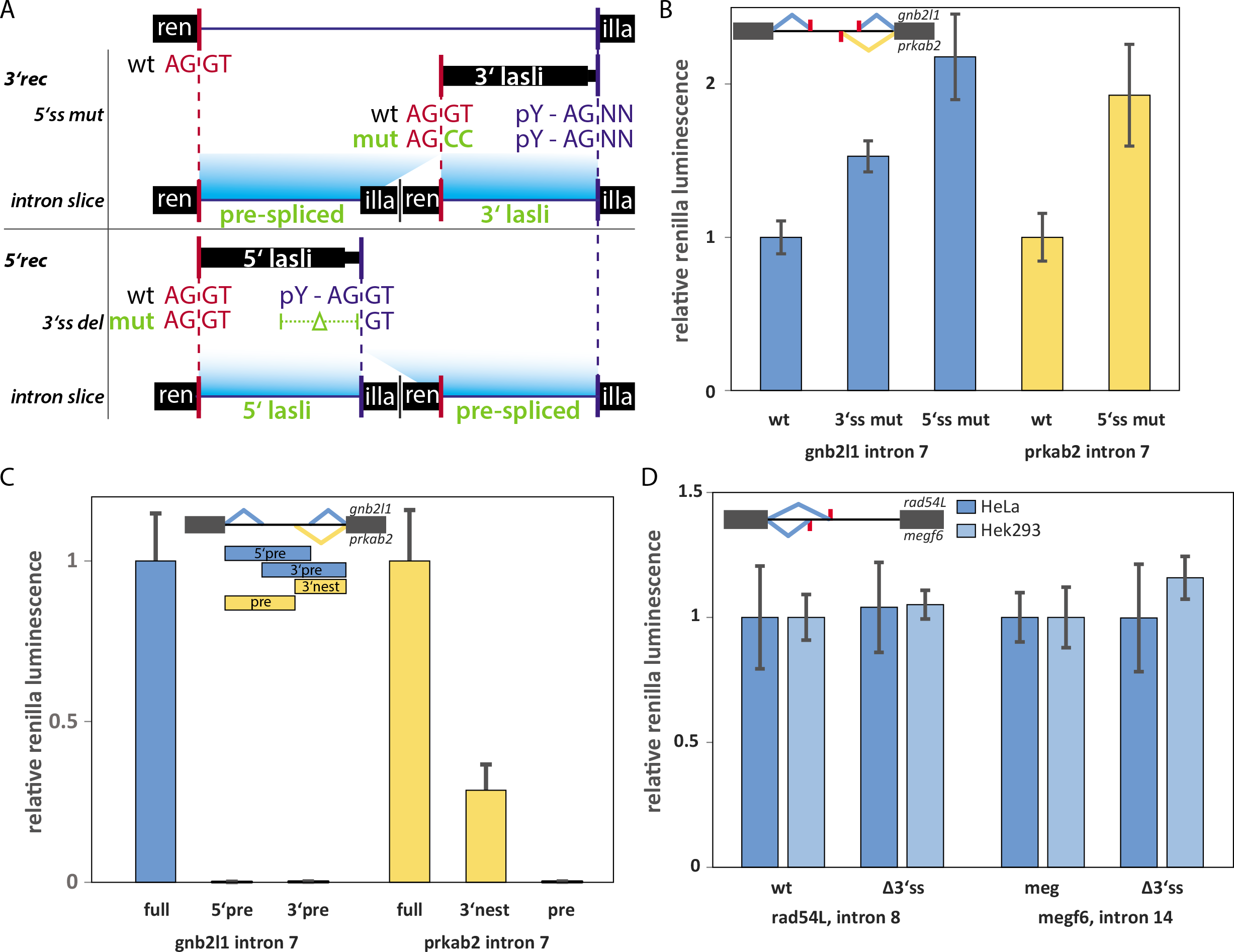
(A) Overview of mutational analysis of intrasplicing event containing introns cloned into the renilla luciferase gene. Putative 3’recursive splice events had their 5’ss mutated and, in some cases, the intron was sliced into the fractions that contain only the intrasplicing event and those that contain the remainder of the intron. These were termed pre-spliced and 3’/5’ lasli, respectively. In case of a 5’ intra splicing event, the 3’ splice site was rendered dysfunctional by deleting the polypyrimidine tract and the downstream AG of the 3’intrasplicesite. (B) Intron 7 of gnb2l1 contains two intrasplicing events. Mutation of the respective intronic splice site leads to an increase in renilla expression (left panels, p<0.05). Similar effects are observed for intron 7 of prkab2, which contains one putative 3’recursive splice event (p<0.05). (C) Prespliced constructs of gnb2l1, intron 7 and prkab2, intron 7 are splicing incompetent and do not produce a functional renilla luciferase (p<0.05). The intronic splicing event of prkab2, intron 7 on its own is well capable of splicing, though apparently with a lower efficiency than the full intron. (D) Example of two genes, rad54L and megf6, where the mutation of the intra-splice sites did not affect full intron removal efficiency (p>0.1).

### Intrasplicing competes with full intron splicing suggesting a novel type of gene expression regulation

The two genes *gnb2l1* and *prkab2* both contained putative recursive splicing events in intron 7. Our dataset included a 5’rec and a 3’rec event for *gnb2l1* and a 3’rec event for *prkab2*. Mutation of these intra-splice sites in our reporter construct increased renilla expression and transcription in all three cases (Figure 7B, Figure S 2), meaning that full intron removal is more efficient in the absence of intrasplicing. Thus, these events are unlikely to be part of a recursive splicing cascade. Removal of the 3’end of intron 7 of both, *gnb2l1* and *prkab2*, does not lead to the reconstitution of a new 3’splice site and the polypoyrimidine tract is removed, rendering the remainder of both introns splicing incompetent. The level of increase of renilla expression by 2-fold for 5’ss mutants allows a rough estimate that 50% of transcripts of this reporter, are spliced at the LaSLi. The inefficient downstream processing of pre-spliced introns is confirmed by three more constructs that only contained the 5’ or 3’ remainder sequence that is left after an initial intra-splicing event. These constructs are unable to express functional luciferase (Figure 7C, “pre”). Contrary, the intrasplicing event of *prkab2*, intron 7 alone is effectively spliced (Figure 7C,”3’nest”). These results are further supported by semiquantitative RT-PCR (Figure S 2). As these splicing events do not facilitate full intron removal, as the model of recursive splicing would suggest, but rather counteract it, they represent, in concert with a downstream degradation mechanism, a potential new mode of regulation of gene expression.

### Silent intrasplicing

We also tested two candidate introns containing putative splicing events at the 5’end, where the pre-splice step does not seem to influence the full intron removal efficiency in the cell lines tested (Figure 7D). This may be either due to this event not taking place in the cell lines we tested, weak and therefor easily outcompeted intra-splice sites or a recursive splicing cascade that is, due to the shortness of the full intron, not a necessity for intron removal.

### Intrasplicing boosts full intron removal

A third type of intrasplicing effect was identified in intron 8 of rbm17, which harbours 3 putative intrasplicing events. Mutation of the intra-splice sites, be it 5’ or 3’ss, led to a strong reduction of renilla expression, both on transcript and protein level (Figure 8A, B). Therefore, these splicing events might be starting points of a recursive splicing cascade, yet due to their positioning, presumably of three different cascades. This is an interesting case, as intron 8 of rbm17 is only slightly over 1kb long and thus, recursive splicing should not be a necessity for this intron’s removal.

**Figure 8:**
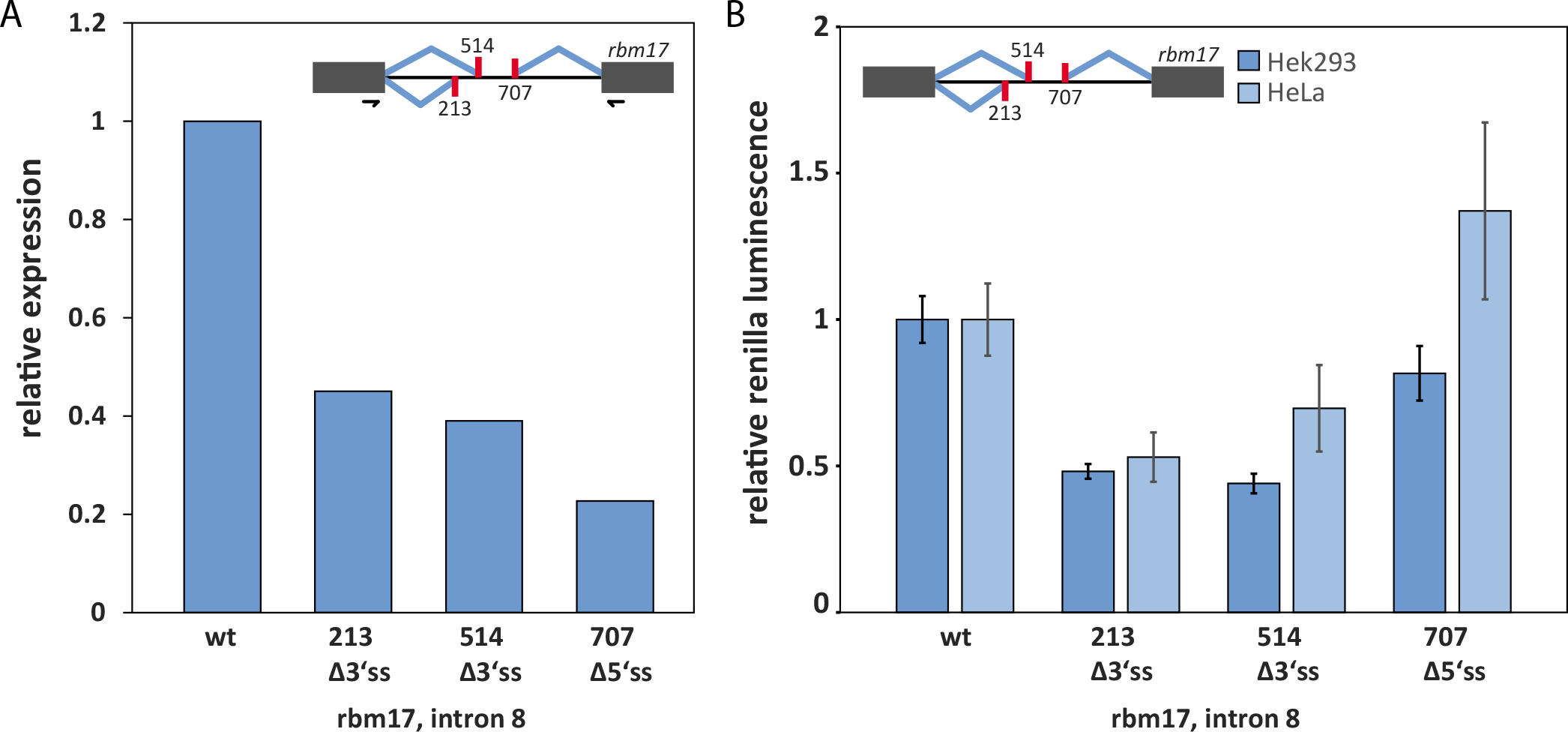
rbm17, intron 8 harbours 3 potential intrasplicing events. Mutation or deletion of the intrasplice sites leads to a strong reduction in transcriptional (A, one representative of two replicates shown) and translational (B) output (p<0.05 for all constructs in Hek cells and 213 in Hela).

### More to discover

Besides the above described modes of intrasplicing, we found several other interesting intrasplicing localizations in our dataset: *epb41l5* expresses several isoforms, two of which are significantly shorter and terminate with an exon that is skipped in the longer isoform. In this terminal intron, we found a recursive splicing event that reconstitutes a RS-site and induces a second splicing step that skips the alternative terminal exon, allowing transcription of the long isoform (Figure 8: rbm17, intron 8 harbours 3 potential intrasplicing events. Mutation or deletion of the intra-splice sites leads to a strong reduction in transcriptional (A, one representative of two replicates shown) and translational (B) output (p<0.05 for all constructs in Hek cells and 213 in Hela). Figure 9A). Attempts to mask this RS-site with antisense oligomers failed to show an isoform shift. Similar has been observed in other studies on recursive splicing, where ASOs were used^9^, either rendering this approach inapplicable or hinting to technical difficulties when targeting ASOs to deep intronic sites.

**Figure 9:**
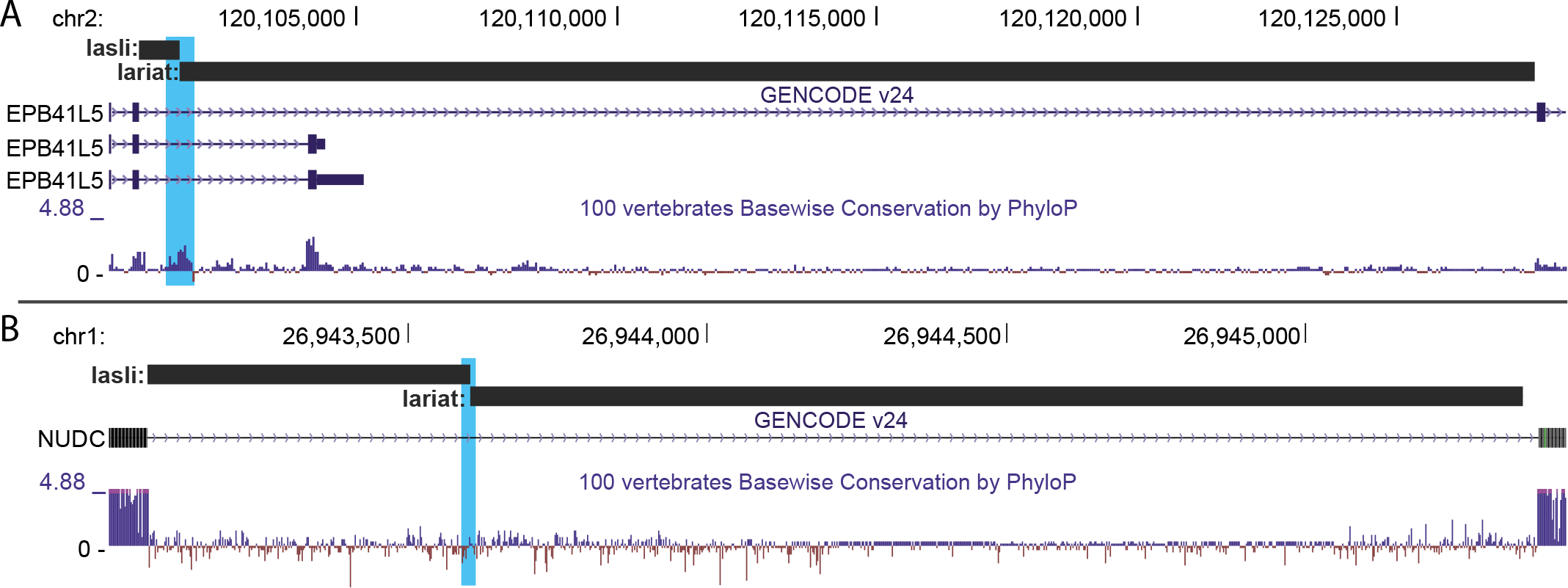
UCSC genome browser screenshots of splicing events. (A) *epb41L5* holds a recursive splicing event in a terminal intron, that induces terminal exon skipping and thus a shift towards expression of a longer isoform. (B) shows a classical recursive splicing situation in a notably short intron (2500bp).

Another example shows two adjacent splicing events in *nudc*. This appears to be a classical recursive splicing situation, yet the intron length is with 2500bp quite short and arguably, would not require recursive splicing as a means of full intron removal.

## Discussion

Currently available gene annotations have a very exon-focused view on the transcriptome. Our splicing dataset aims at extending available annotations and promoting a more processing-aware approach to transcription. Generation of mini-genes or mini-introns as well as deep intronic mutations can result in unprocessed or inefficiently transcribed RNAs due to insufficient understanding of intronic elements controlling RNA processing and thus stability. In combination with other available datasets or tools such as eCLIP-seq^26^, splicePie^17^ or computational prediction of regulatory sequence elements by tools like RBPmap^27^, a broader understanding of the processing steps required to achieve regulated gene expression can be obtained.

Current research, dealing with splice junction discovery on a massive scale^28^ clearly showed that splicing cascades are by far more complex than current annotations imply. Yet, the extensiveness of this dataset makes it close to impossible to determine biological relevance of individual splice reactions. In our study we attempted, with a more restrictive approach, to extract high confidence events that can be more easily tested and we evaluated their impact on gene expression. Our pipeline utilizes tools designed for the discovery of circular RNAs to supplement the information a regular split read discovery gains on splice site location. Besides increasing confidence in the discovered splice junctions, this also allows for the localization of the branchpoint. Even though we only took a closer look at a handful of unusual splicing events, our results anticipate a much broader range of impacts of intrasplicing events on gene expression. Based on our data and on the diversity of mechanisms we found, we can speculate on the diverse effects of intrasplicing events on full intron splicing. Recursive splicing allows for clean full intron removal. Yet, the necessity of RS-sites for efficient full intron removal remains debated. In some cases, recursive splicing might be a means of processing a mis-spliced intron, to retain transcript functionality. This might be especially true for short introns, where recursive splicing seems not be an approach to spanning long distances between splice sites. Mis-splicing could be promoted by an abundance of U1-binding in many introns, as premature cleavage and polyadenylation (PCPA) inhibitor^7^, what might direct the splicing machinery to falsely recognize intronic splice sites and commit to an intrasplicing event.

If the transcript cannot be further processed or processed in a way that does not interrupt the reading frame, situations like in *prkab2*, intron 7 splicing can occur. Here the intrasplicing event effectively removes the functional 3’splice site, branchpoint and polypyrimidine tract, rendering the remainder of the intron retained (Figure 10A). Induction of such a dead-end processing in combination with downstream degradation pathways, such as nonsense-mediated decay, can be a powerful regulatory tool for gene regulation. It also lays the foundation for exon skipping induction, as depicted in Figure 10B. Vice versa, speeding up processing of dwelling introns can have the opposite effect by overcoming nuclear retention and facilitating mRNA export. Even with the precise mechanism unknown, such a situation seems to occur in *rbm17*, intron 8, presumably via recursive splicing. Yet unknown is the role of the exon junction complex (EJC) in recursive and intrasplicing. CliP experiments determining genome-wide EJC deposition^29^ clearly show that our intrasplicing events are associated with proximal intronic EJC binding (Figure S 3). The consequences of EJC deposition are broad and range from modulating splicing patterns, transcription speed, mRNA stability to export and translation efficiency. The effect of EJC deposition on introns has not been studied to date but it seems likely that interference with spliceosome assembly would occur. In order to effectively splice the full intron in recursive or intrasplicing patterns, this interference has to be handled. Therefore, further studies are required to elucidate the role of EJCs in recursive and intrasplicing.

**Figure 10:**
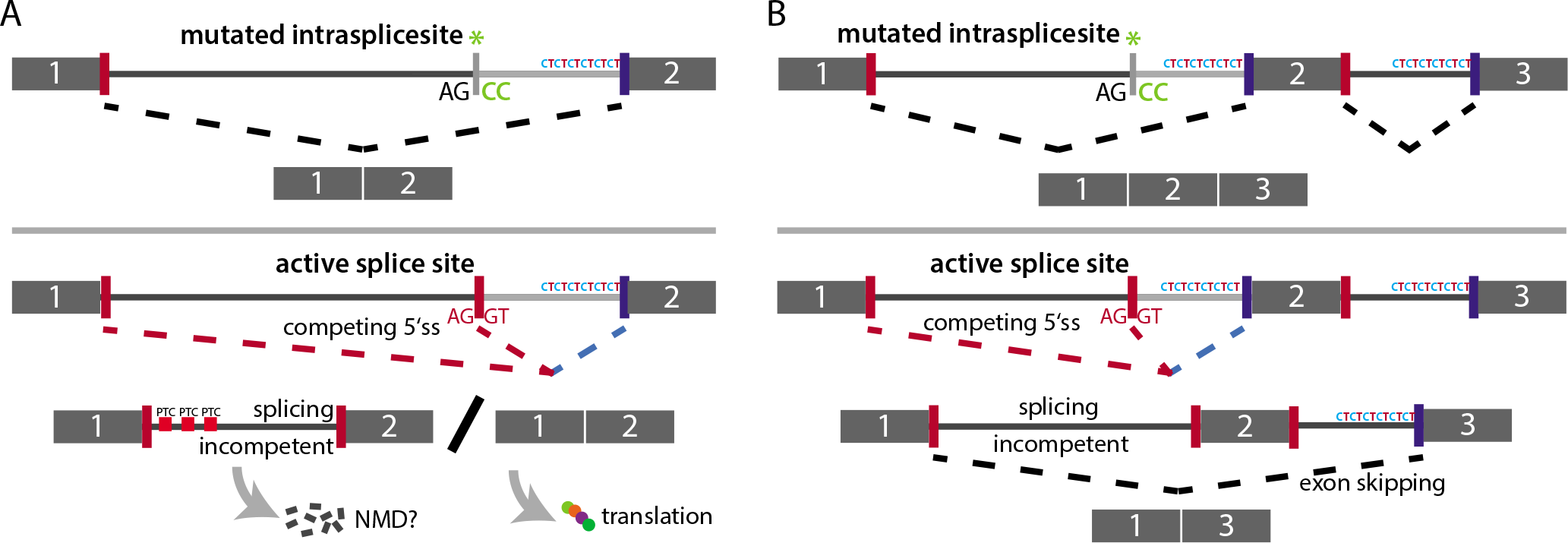
Two models illustrating the potential impact intrasplicing events can have on gene expression, via regulation of transcript abundance and isoform expression by inducing exon skipping events. (A) In absence of a functional intra-splice site, the intron is efficiently spliced. An intrasplicing event on the other hand removes the py-tract and does not re-establish and RS-site. As a result, the remainder of the intron is retained, leading to nuclear retention or NMD. (B) If a downstream intron is present, such an intrasplicing event can induce exon skipping by utilizing the next functional 3’ss.

Recent studies utilizing nascent RNA sequencing to determine the co-transcriptional nature of splicing have shown that intron removal is in many cases a prompt event and occurs seconds after the splice sites have been transcribed^30,31^. If this observation can be extended to human transcription, it explains high occurrence of intrasplicing we observe, despite partially weak splice site interactions (Figure S 1).

The variety of mechanistical implications makes intrasplicing a promising new regulatory layer of gene expression that adds to the already complex and dynamic regulatory landscape of multi-exon gene regulation. Future studies will focus on the interactors involved in intra-splice site recognition and a potential cell-type specific cocktail of splicing factors that allows fine tuning of the transcriptome via intra-splicing.

## Materials and Methods

### Cell culture and transfection

HeLa and Hek293T cells have been grown in DMEM medium, supplemented with 10% FCS (Sigma-Aldrich). Cells were transfected with Lipofectamine 3000 (Thermo Fisher) according to the manufacturer’s specifications.

### RNA preparation, reverse transcription and PCR

RNA was isolated 48 hours post-transfection with Tri-reagent (Sigma-Aldrich), DNAse I (Roche) treated with 2x 20U per 50µg RNA for 30 minutes and then phenol chloroform extracted. Reverse transcription was carried out with superscript III (Invitrogen) on 1 µg total RNA for 60 minutes at 50°C. 1/20th of this reaction was then used as a template for subsequent PCR (oneTaq, NEB) or qPCR (Evagreen, Medibena). Lariat PCR was carried out with diverging primers, spanning the putative branch point. In some cases, nested PCR with cycle numbers ranging from 20 to 30 cycles was necessary to amplify lowly abundant lariats. All primers used for PCR, qPCR and cloning can be found in the supplementary data.

### Cloning and site directed mutagenesis

The phRL-TK constructs containing various introns have been generated with the InFusion cloning kit (Clontech) and NEBuilder (NEB). All introns have been inserted at position 414 of the renilla gene. The splice site mutations and nested intron deletions have been introduced with the Q5 site-directed mutagenesis kit (NEB).

### Dual luciferase assay

For the luciferase assay, the experimental phRL-TK vector with intron insertions has been cotransfected with pmirGLO –ren as an internal control in 24-well plates. pmirGLO −ren is a modified version of pmirGLO, where the renilla gene and its promoter have been removed by restriction digest and religation. 48 hours after transfection the medium, except 100 µL was removed and the luciferase assay was carried out according to the manufacturer’s specifications. Luminescence was measured on a luminometer (Robion Solaris 3170) and relative renilla luminescence was calculated by background subtraction and normalization to the internal firefly control.

### Bioinformatic pipeline for splicing event extraction

The focus on pre-mRNA and lariats implied the usage of datasets that are potentially enriched in these RNA species. Therefore the main focus of this approach was on two RNA-seq datasets: nuclear, non-ployadenylated long RNA and RNAse R digested circular RNA. The raw RNA sequencing data of these datasets, together with several others, to extend the read pool (Table 1), were obtained from the gene expression omnibus (https://www.ncbi.nlm.nih.gov/geo) and have been processed by the following pipeline:

a) quality control: adapter clipping, quality trimming and duplicate removal where necessary with fqtrim (http://ccb.jhu.edu/software/fqtrim/index.shtml) and FastUniq (https://sourceforge.net/projects/fastuniq/).
b) Mapping to human reference genome hg38 with segemehl 0.1.9^18^. The segemehl tool has been modified to allow detection of split reads that are up to 1.2 million nucleotides apart, which ensures detection of split reads covering the largest introns in the human genome.
c) For the extraction of split reads, segemehl’s realign routine was run. The resulting bedfile * .splice.bed was processed with bedtools^32^ and custom scripts, extracting and sorting linear and circular split reads and filtering those out that did not overlap with annotated introns and extended beyond gene boundaries.
d) In order to obtain high confidence splicing events, intronic circular and linear splits were intersected and filtered by following requirements: the 5’ ends of intersecting linear and circular splits cannot deviate more than 2 nucleotides in any direction, to tolerate a given uncertainty in split-read mapping; the 3’ end of the circular split has to be located within 100 nucleotides upstream of the 3’ end of the linear split read to allow for a certain range of branchpoint-splice site distances.

**Table 1:** datasets accessible via GEO or SRA, used for splicing event extraction.

| Dataset | GEO/SRA accession | Library preparation | reference |
| --- | --- | --- | --- |
| 12tissues | GSE45326 | total RNA | Nielsen et al., 2014 <sup>35</sup> |
| Illumina body map 2.0 | GSE30611 | total RNA | http://www.ncbi.nlm.nih.gov/geo/query/acc.cgi?acc=GSM759490 |
| Brain single cell | GSE67835 | total RNA, single cell | Darmanis et al., 2015 <sup>36</sup> |
| **circSeq** | SRP011042 | total RNA, RNase R | Jeck et al., 2013 <sup>20</sup> |
| **polyA minus** | GSE22666 | polyA minus RNA | Yang et al., 2011<sup>19</sup> |
| **BPseq** | GSE53328 | lariatSequencing + BPCapture | Mercer et al., 2015<sup>16</sup> |

It is not possible to precisely allocate lariats and their respective linear split if, for example an alternative 3’ splice site is used and two branchpoints and therefore 3’ ends of two different circular splits lie within 100 nucleotides upstream of this splice site. This results in a small number of redundant splicing events, varying in BP and 3’ss position.

The resulting splicing events of this extraction procedure were termed lariat-spanning linear splits (LaSLiS) and provide the basis of all subsequent analyses.

### Splicing event analyses

LaSLiS were classified based on the position within the hosting intron. LaSLiS with exon contact and therefore utilizing one primary splice site were classified as potentially recursive and LaSLis without any exon contact as intra- or nested splicing events.

Sequences of LaSLiS splice sites (i.e. the 5’ and 3’ ends and surrounding nucleotides) were extracted and motifs were generated with Weblogo 3^33^. As a reference, 5’ and 3’ ends of annotated introns were analyzed with the same procedure.

Conservation scores for seven vertebrates were obtained from the phyloP7 project^34^ (http://hgdownload.cse.ucsc.edu/goldenPath/hg38/phyloP7way/) and intersected with the 20 nucleotides surrounding the 5’ and 3’ ends of LaSLiS. The reference is another window of 20 nucleotides from within every intron (>500 nts) in the human genome.

Splice site scoring was achieved with Xmaxentscore^21^. Respective sequences were extracted and used as input. If, due to poor split-read mapping, a higher splice site score was found within +/- 2 nts of the 5’ or 3’ end of any LaSli, the higher scoring position was assumed as the proper splice site. This was done to increase fidelity in splice site annotation. Both, the original and the optimized dataset are available.

## Figure Legends

**Figure S 1:**
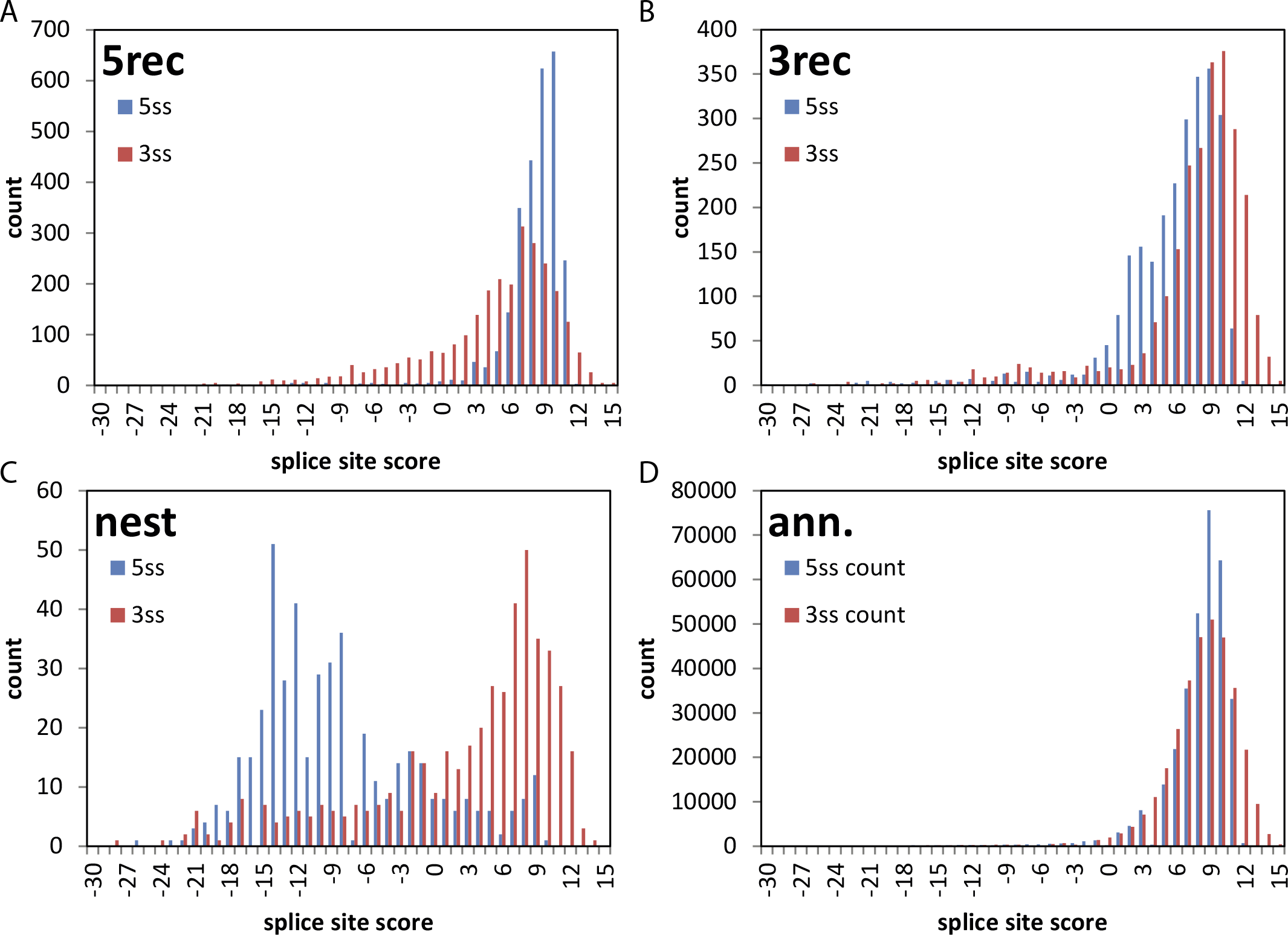
Maximum entropy score distribution for each given class of splice sites. The computed score is given on the x-axis and the number of splicing events in each given subset is shown on the y-axis. (D) shows the scores of annotated splice sites and is used as a reference set.

**Figure S 2:**
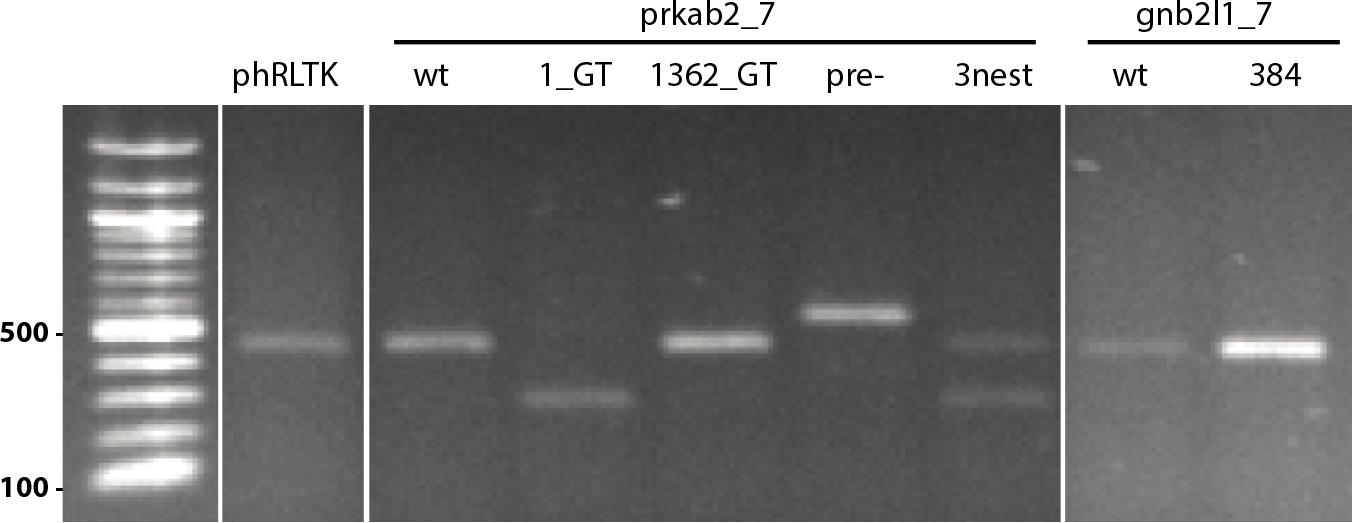
RT-PCR of prkab2 and gnb2l1 intron 7 in phRL-TK including splice site mutations. This semi-quantitative PCR reproduces the result of the renilla assay and sheds additional light on RNA processing. A mutation of the 5’ss of prkab2, intron 7 leads to alternative splice site selection within the renilla exon. This splice site is also used when only the 3’intrasplicing event is cloned into the reporter. Here about 50% of transcripts use the native, the rest the exonic splice site. This reflects the reduced intra-splice site strength. The longer product of the pre-spliced transcript arises due to the incomplete processing and splicing incompetence.

**Figure S 3:**
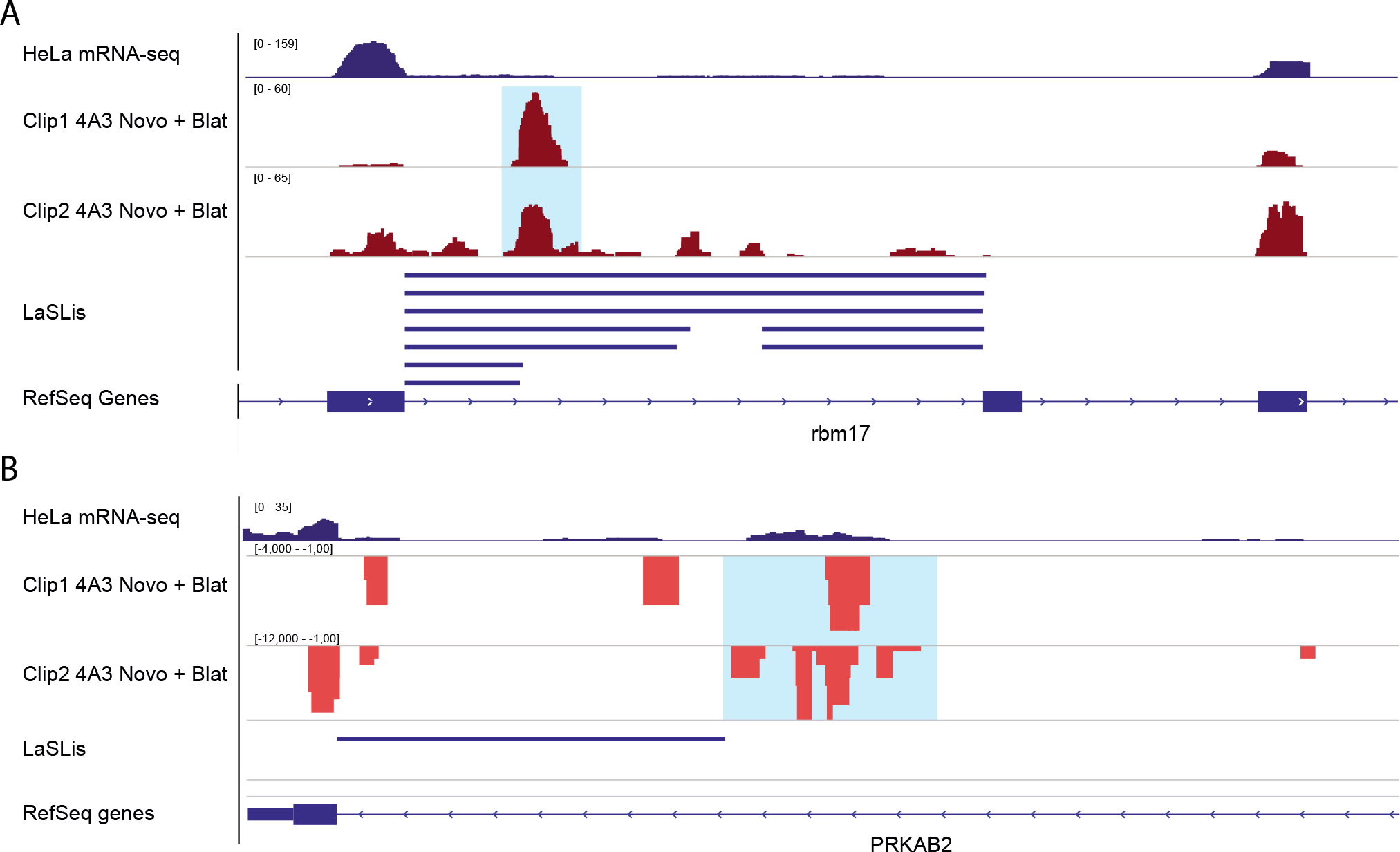
Exon junction clip data from Saulière et al. (2012)^29^ shows clear deposition of exon junction complexes in genomic regions flanking intrasplicing events (blue boxes).

